# Double drives and private alleles for localised population genetic control

**DOI:** 10.1101/2021.01.08.425856

**Authors:** Katie Willis, Austin Burt

**Author notes:** Corresponding author, (KW).

## Abstract

Synthetic gene drive constructs could, in principle, provide the basis for highly efficient interventions to control disease vectors and other pest species. This efficiency derives in part from leveraging natural processes of dispersal and gene flow to spread the construct and its impacts from one population to another. However, sometimes (for example, with invasive species) only specific populations are in need of control, and impacts on non-target populations would be undesirable. Many gene drive designs use nucleases that recognise and cleave specific genomic sequences, and one way to restrict their spread would be to exploit sequence differences between target and non-target populations. In this paper we propose and model a series of low threshold double drive designs for population suppression, each consisting of two constructs, one imposing a reproductive load on the population and the other inserted into a differentiated locus and controlling the drive of the first. Simple deterministic, discrete-generation computer simulations are used to assess the alternative designs. We find that the simplest double drive designs are significantly more robust to pre-existing cleavage resistance at the differentiated locus than single drive designs, and that more complex designs incorporating sex ratio distortion can be more efficient still, even allowing for successful control when the differentiated locus is neutral and there is up to 50% pre-existing resistance in the target population. Similar designs can also be used for population replacement, with similar benefits. A population genomic analysis of PAM sites in island and mainland populations of the malaria mosquito *Anopheles gambiae* indicates that the differentiation needed for our methods to work can exist in nature. Double drives should be considered when efficient but localised population genetic control is needed and there is some genetic differentiation between target and non-target populations.

**Author summary:** Some disease vectors, invasive species, and other pests cannot be satisfactorily controlled with existing interventions, and new methods are required. Synthetic gene drive systems that are able to spread though populations because they are inherited at a greater-than-Mendelian rate have the potential to form the basis for new, highly efficient pest control measures. The most efficient such strategies use natural gene flow to spread a construct throughout a species’ range, but if control is only desired in a particular location then these approaches may not be appropriate. As some of the most promising gene drive designs use nucleases to target specific DNA sequences, it ought to be possible to exploit sequence differences between target and non-target populations to restrict the spread and impact of a gene drive. In this paper we propose using two-construct “double drive” designs that exploit pre-existing sequence differences between target and non-target populations. Our approaches maintain the efficiencies associated with only small release rates being needed and can work if the differentiated locus is selectively neutral and if the differentiation is far from complete, and therefore expand the range of options to be considered in developing genetic approaches to control pest species.

## Introduction

Gene drive is a natural phenomenon in which some genes are able to increase in frequency and spread through populations by contriving to be inherited at a greater-than-Mendelian rate [1, 2]. Strong drive can cause genes to increase rapidly in frequency even if they also harm the organisms carrying them, and there is currently much effort trying to develop synthetic gene drive constructs (or gene drives) to control disease-transmitting mosquitoes and other pest populations that have thus far been difficult or impossible to manage satisfactorily [3–6]. If a species is harmful and subject to control measures wherever it exists, then, in principle (i.e., in the computer), highly efficient gene drive strategies can be devised that exploit natural processes of dispersal and gene flow such that relatively small inoculative releases in a few locations can lead to substantial and widespread impacts over subsequent generations [7–9]. However, some species are pests only in a part of their range (e.g., invasive species), and other approaches are needed.

Two broad approaches have been proposed for restricting the impact of genetic control interventions to a target population. First, one can use a strategy requiring relatively large releases, which can be restricted to the target population, with any introductions into the non-target population (by dispersal, or by accidental or unauthorised releases) being too small to have a significant impact. Potentially suitable genetic constructs include those that do not drive (e.g., dominant lethals, autosomal X-shredders, or Y-linked editors; [10–12]), or those that do, but only if they are above some threshold frequency (e.g., many approaches based on the logic of toxins and antidotes; [13–15]). Some of these approaches are more efficient than others [10, 16, 17], but, by necessity, all of them require a non-trivial production and release effort.

Alternatively, one could exploit sequence differences between target and non-target populations, in which case it may be possible to retain the small release rates and overall efficiency of low threshold gene drive approaches [18, 19]. Sudweeks et al. [19] present useful modelling of this approach, considering the case where there is a locally fixed allele of an essential gene in the target population, while non-target populations carry a resistant allele at some frequency. A nuclease-based gene drive that uses the homing reaction (i.e., sequence-specific cleavage followed by homologous repair [3, 20]) to disrupt the locally fixed allele could be released into and eliminate the target population, but have little impact, or only a transient impact, on non-target populations. However, as emphasised by the authors, if the target population has even a small frequency of the resistant allele, then that allele could be rapidly selected for and the intervention fail.

In this paper we explore alternative two locus “double drive” low threshold strategies to restrict population control based on pre-existing sequence differences between target and non-target populations. All our designs are based on a division of labour between the two constructs, with one imposing a reproductive load by disrupting a gene needed for survival or reproduction, and therefore responsible for the desired impact (population suppression), and the other responsible for the population restriction. These designs are substantially less susceptible to pre-existing resistance in the target population at the differentiated locus than single drive designs, and can even work if the differentiated locus is selectively neutral. Double drives may also be useful for population replacement. Finally, analyses of published genome sequences from island and mainland populations of the malaria mosquito *Anopheles gambiae* indicates that the sort of population differentiation we model can exist in nature.

## Results

### Simple double drives for population suppression

The simplest double drive designs we consider consist of one construct (call it α) inserted into and disrupting a haplo-sufficient female-essential gene, such that homozygous females die without reproducing while heterozygous females and all males are unaffected, and a second construct (β) inserted into a sequence that is significantly more common in the target than the non-target population(s). Both constructs are able to drive by the homing reaction but α can home only in the presence of β, while β may either home autonomously or rely on the presence of α. With CRISPR-based designs, α would encode its cognate gRNA, β would encode the Cas9, and either construct could encode the gRNA for the second locus (Fig 1, Designs 1 and 2). We assume the α construct has been designed such that functional resistance is not possible, though non-functional resistant alleles can arise by end-joining repair [21]. For the β construct we initially suppose its insertion site (i.e., the differentiated locus) is selectively neutral and unlinked with the α insertion site, and that differentiation is nearly complete, with the target sequence present at a frequency of 99% in the target population and absent in the non-target population (i.e., it is a virtually fixed private allele).

**Fig 1.**
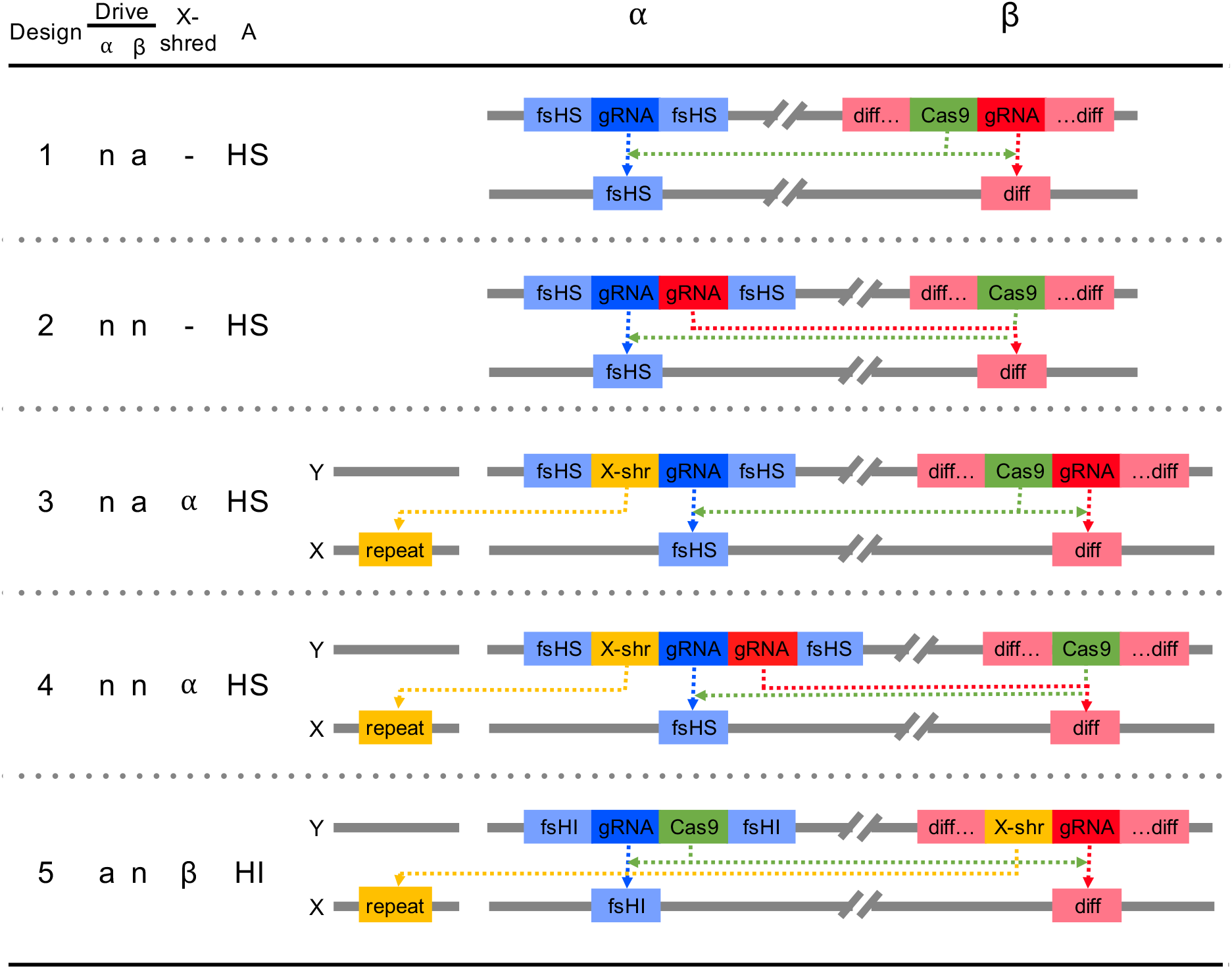
Alternative double drive designs for population suppression. Constructs α and β can drive autonomously (a) or non-autonomously (n); one or the other may encode an X-shredder; and the A target locus can be a gene that is haplo-sufficient (HS) or haplo-insufficient (HI) for female viability or fertility. fsHS - female-specific haplo-sufficient locus; fsHI - female-specific haplo-insufficient locus; diff - differentiated sequence; X-shr - X-shredder targeting an X-linked repeat.

Under these conditions, a small (0.1%) release of males carrying Design 1 constructs into the target population leads to both constructs rapidly increasing in frequency and, as a result, the population size crashes to a minimum size of 3.58e-6 (relative to the pre-release equilibrium) in 25 generations (Fig 2A). Depending on the initial population size and the biology of the species (e.g., whether there are Allee effects [22]), this decline could be enough to eliminate the population. However, in our simple deterministic model population elimination is not possible. Instead, the population recovers due to the evolution of resistance at the differentiated locus, leading to loss of the β construct, which then leads to loss of α, allowing the wild-type allele and population fertility to recover. By contrast, the same releases into the non-target population have minimal effect: β cannot increase in frequency, both construct frequencies remain low, and population size is little affected (Fig 2B).

**Fig 2.**
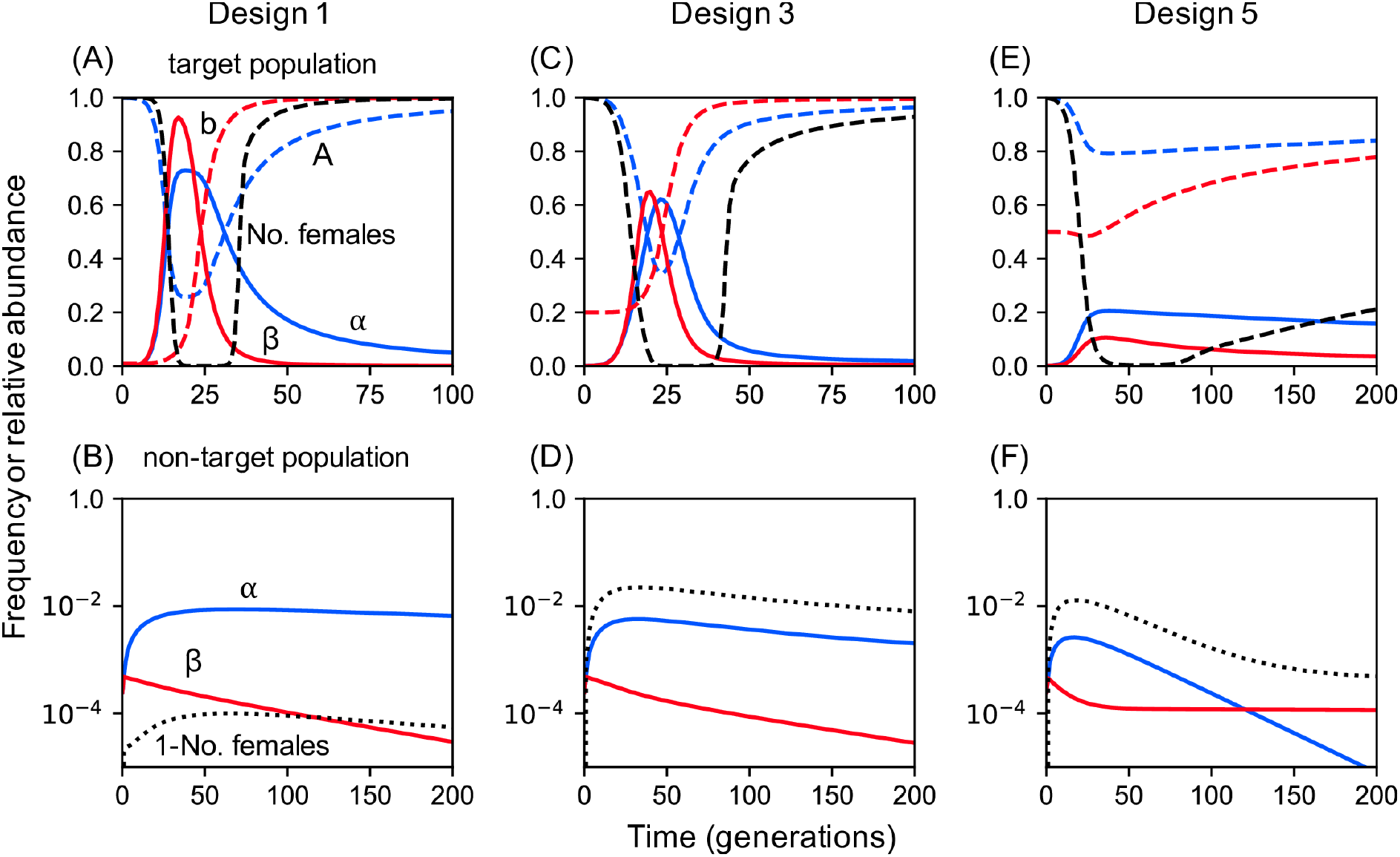
Performance of double drives for population suppression. **(A, B)** Timecourse for Design 1 in target and non-target populations, assuming 1% and 100% pre-existing resistance at the B locus, respectively. In the target population the α and β constructs increase in frequency together (blue and red solid lines), causing the number of females to decline. If the population is not eliminated, then eventually the resistant b allele replaces β, followed by the wild-type A allele replacing α, allowing the population to recover. In the non-target population both constructs remain rare and the reduction in female numbers remains small. **(C, D)** Timecourse for Design 3 assuming 20% and 100% pre-existing resistance in the target and non-target populations. **(E, F)** Timecourse for Design 5 assuming 50% and 100% pre-existing resistance in the target and non-target populations.

Because the spread of construct α in the target population depends on β, and therefore will be affected by the association between them, it might be expected that close linkage between the two constructs may increase construct spread and the extent of population suppression. Furthermore, because the population may eventually recover due to the evolution of resistance at the differentiated locus, additional improvements might be expected by using an essential gene as the differentiated locus and designing β to have minimal fitness effects (e.g., by containing a recoded version of the target gene [23–25], or being inserted in an artificial intron [26]). With such a design end-joining repair will tend to produce non-functional resistance alleles, and resistance will be slower to evolve, relying instead on pre-existing resistant alleles. Both these expectations about linkage and using an essential differentiated gene are met individually, and, in combination, can reduce the minimum population size achieved by many orders of magnitude (Fig 3; see also S1 Fig for the separate effect of each modification). If it is not possible to have close linkage, then the maximum level of suppression can also be increased by releasing the two constructs in different males rather than in the same males, though at the cost of the impact being delayed, and separate releases perform worse than combined releases when linkage is tight (S2 Fig).

**Fig 3.**
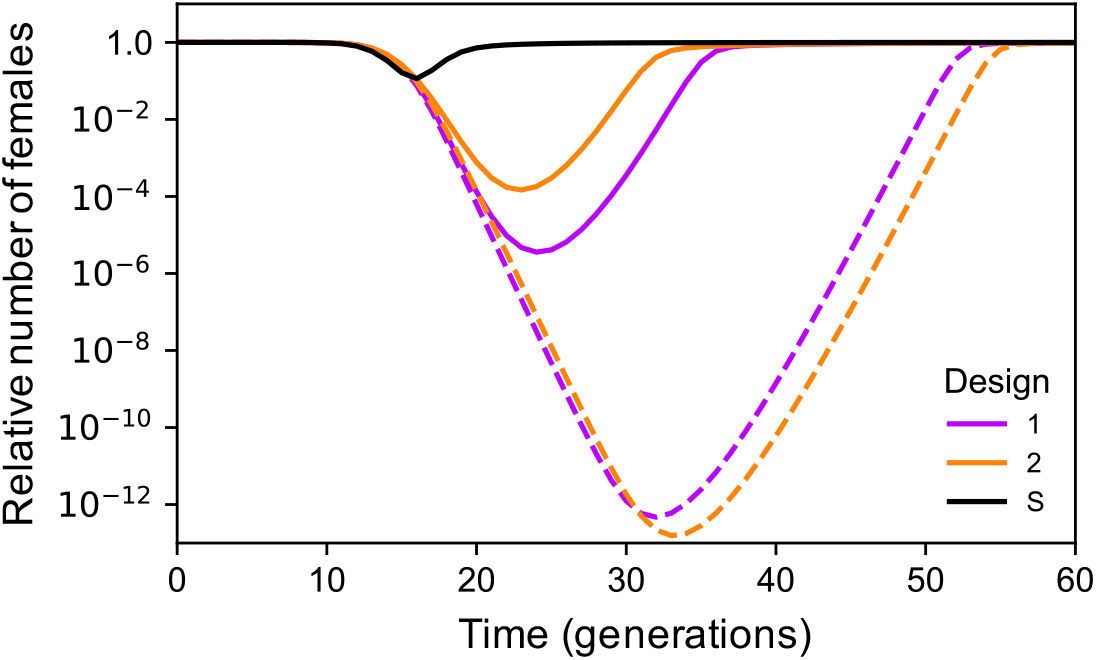
Timecourse for the relative number of females over time for Designs 1 and 2. Solid lines are for β in a neutral locus unlinked to the *α* construct, and dashed lines for β as a neutral insertion in a haplo-insufficient essential gene closely linked (r=0.01) to the *α* construct. In all cases there is 1% pre-existing resistance at the B locus. Also shown for comparison are results for a single construct drive targeting a haplo-sufficient female-specific viability gene (S).

Design 2, which has the same components as Design 1, but arranged differently such that homing of the β construct only occurs in the presence of α, has dynamics qualitatively similar to Design 1, but quantitatively different (S3 Fig). Interestingly, if the two constructs are unlinked then the extent of suppression is less than with Design 1, but if they are closely linked then the suppression can be greater (Fig 3 and S1 Fig). All these possible alternatives give significantly greater suppression than a single drive imposing a reproductive load by targeting a female-essential gene (Fig 3).

### Coping with higher frequencies of pre-existing resistance

Though these simple double drive designs work well with pre-existing target site resistance at the differentiated locus of 1%, performance declines rapidly after that. For example, if there is 10% pre-existing resistance, then even the best of these designs (Design 2 with close linkage and the differentiated locus being haplo-insufficient) only suppresses the target population to a minimum of 2.38e-4. In some situations the target population may not have a private allele with frequency over 90% and alternative approaches would need to be considered. One possibility is to increase the load imposed on the population by the α construct by adding to it an X-shredder locus that destroys the X-chromosome during spermatogenesis such that it produces a male-biased sex ratio as well as killing heterozygous females (Fig 1, Designs 3 and 4). Since population productivity in many species depends on the number of females, population size may thereby be further reduced. A single drive based on these components has previously been constructed in *Anopheles gambiae* by Simoni et al. [27]. Our modelling indicates that adding an X-shredder to a double drive gives a quantitative improvement in the dynamics, and even pre-existing resistance frequencies of 20% are compatible with good control, while still having minimal effect on non-target populations (Fig 2C and D).

Even more robust control can be obtained by adding the X-shredder to the β locus and having the α locus drive autonomously in males and cause dominant sterility or lethality in females (e.g., target a female-specific haplo-insufficient locus; Fig 1, Design 5). The dynamics in this case are somewhat different from the others: the X-shredder does not function to directly increase the load, but instead it allows the α construct to spread in the population, because it will end up more often in males (where it homes), and less often in females (where it is a dead end). The male bias also protects the β construct from the female lethality produced by the α construct, and so selection against β is much weaker than in the previous designs, and resistance evolves more slowly (compare the rate of spread of the resistant b allele in Fig 2E to that in Fig 2A and C). As a result the design is able to perform well even with pre-existing resistance of up to 50%, but still not spread in the non-target population (Fig 2E and F). Moreover, if the population is not eliminated, it can nevertheless be suppressed for many generations. For example, with 50% pre-existing resistance the minimum population size reached is 2.15e-3, and the population remains below 5% of its pre-intervention size for 63 generations; with close linkage (r=0.01), then the corresponding values are 8.79e-7 and 147 generations. A comparison of the maximum extent of suppression as a function of the pre-existing resistance frequency for the different designs is shown in Fig 4 (see also S4 Fig). With all the modifications considered (linkage, use of an essential differentiated gene, or separate releases) effects on the non-target population remain small (S5 Fig).

**Fig 4.**
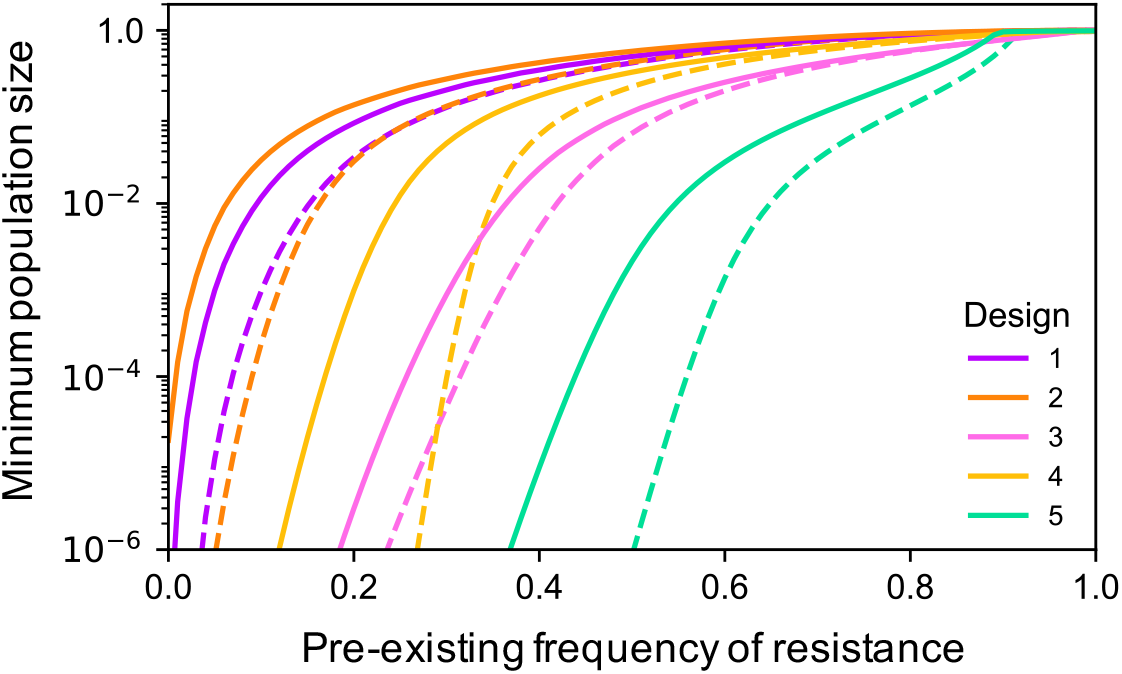
Minimum population size for each of the 5 designs as a function of the pre-existing frequency of resistance. Solid lines are for the baseline case (r=0.5, β in a neutral locus), while dashed lines are for the improved case (r=0.01, β as a neutral insertion in a haplo-insufficient essential gene).

### Evolutionary stability and impact of fitness costs

We now explore the consequences of relaxing two assumptions that have been implicit thus far in our modeling. First, we have assumed that our various constructs remain intact after release. In fact, mutations that destroy the function of one component or another will be expected to arise as the constructs spread through a population, particularly as homing may be associated with a higher mutation rate than normal DNA replication [28]. For components that contribute directly to their construct’s spread, one would expect that loss-of-function mutations would remain rare in the population and have little effect, whereas for other components (e.g., the X-shredder), such mutations may be actively selected for. To investigate we allowed homing-associated loss-of-function mutations to occur in each component of each construct. Mutation rates of 10e-3 have a small but significant impact on the performance of the three designs with an X-shredder, due to the accumulation of mutant constructs missing that component, while mutation rates of 10e-4 have negligible impact for all designs (Fig 5A, S6 Fig and S7 Fig).

**Fig 5.**
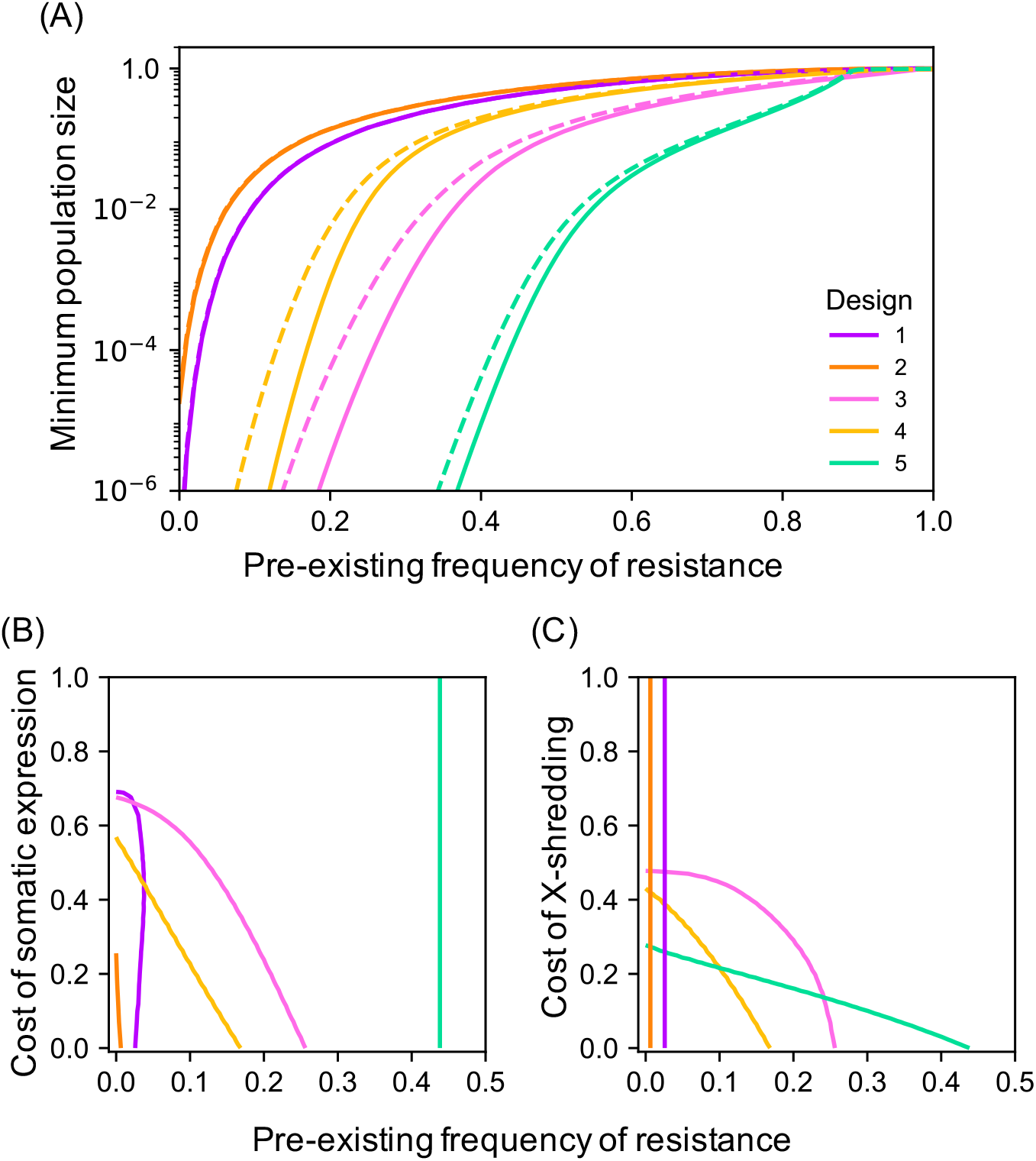
The impact of evolutionary stability and added fitness costs on the performance of each of the 5 designs for population suppression. **(A)** The effect of loss-of-function mutations on the minimum population size reached. Solid lines are for the baseline case of no mutations, and dashed lines are with each component of each construct having a mutation rate of 10e-3 per homing event. Note that the effect is only visible for designs with an X-shredder, and if the mutation rate was 10e-4, the results for all designs would be virtually indistinguishable from the solid lines. **(B, C)** Contour plots showing combinations of fitness costs and pre-existing frequency of resistance giving a minimum population size of 10e-4 for different double drive designs. For **(B)** the costs are reductions in female fitness due to somatic expression of the nuclease targeting the A locus, and for **(C)** the costs are reductions in male fitness due to the X-shredder. Vertical lines indicate the cost is irrelevant, either because heterozygous females in any case have fitness 0 (Design 5 in **(B)**), or because the designs do not include an X-shredder (Designs 1 and 2 in **(C)**).

Second, we have assumed thus far that the genetic constructs have little unintended impact on survival or reproduction. Experiments with *An. gambiae* have revealed at least two unintended fitness costs can occur, a reduced fitness of homing heterozygous females due to somatic expression of the nuclease [21, 27], and reduced fitness of males expressing an X-shredder, possibly due to paternal deposition of the nuclease and/or reduced sperm production [29]. The first of these costs is not relevant to Design 5 (because heterozygous females die anyway), and the second is not relevant to Designs 1 and 2 (because they do not use an X-shredder), but in other contexts, as expected, these costs reduce performance, requiring a lower frequency of pre-existing resistance in order to achieve a particular level of suppression (Fig 5B and C).

### Population replacement

Gene drive can be used not only for population suppression but also to introduce a new desirable ‘cargo’ gene into a target population for population replacement or modification for example, a gene reducing a mosquito’s ability to transmit a pathogen [30, 31]. In double drive designs for population replacement the α construct would carry the cargo and homing by α would require β, while that by β could be either autonomous or depend on α (Fig 6A). Both α and β could be inserted into neutral sites, or into essential genes in such a way as to minimise fitness effects. We have modeled these approaches assuming, for purposes of illustration, the cargo imposes a dominant 20% fitness cost on females, and find that, again, such double drives can spread rapidly through target populations even when there is significant pre-existing resistance, and would not spread in non-target populations fixed for the resistant allele (Fig 6B and C). Unless there is virtually no pre-existing resistance at the differentiated locus, double drives can keep the frequency of the cargo gene above 95% much longer than a single drive construct targeting a differentiated locus, either neutral or essential (Fig 6D). If α is inserted in an essential gene, protection can be even longer.

**Fig 6.**
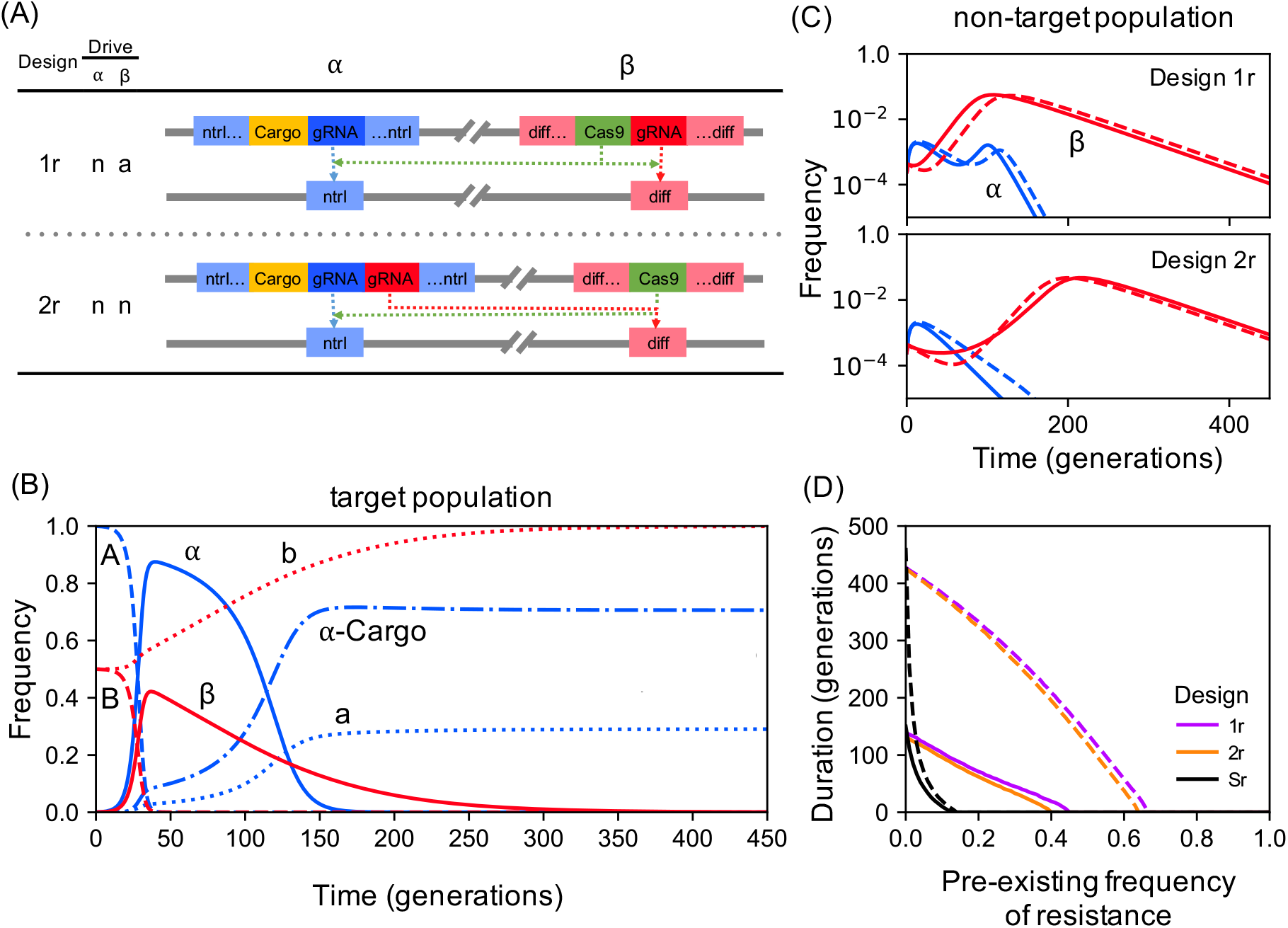
Double drives for population replacement. **(A)** Alternative double drive designs. **(B)** Timecourse of allele frequencies for Design 2r in a target population assuming 50% pre-existing resistance. The dynamics for Design 1r are qualitatively similar. **(C)** Allele frequencies for Designs 1r and 2r in a non-target population with 90% pre-existing resistance, assuming insertion of both *α* and β into neutral loci (solid lines); or *α* as a neutral insertion into a haplo-insufficient essential gene (dashed lines). **(D)** Duration of at least 95% of adult females carrying the cargo-bearing *α* construct as a function of the pre-existing frequency of resistance at the B locus for double drives 1r and 2r, where solid and dashed lines are as in **(C)**. Also shown for comparison are results for a single drive (S) carrying the cargo at a neutral A locus (solid line) or a haplo-insufficient essential gene (dashed lines). Note that results for insertion of β into a haplo-insufficient essential gene would be virtually indistinguishable from the solid lines **(C, D)**. All plots assume 20% fitness cost of the cargo on females and a homing-associated loss-of-function mutation rate of 10e-3.

### PAM site analysis in An. gambiae

To explore whether the type of population differentiation assumed in our modelling can exist in nature, we analysed published genome sequence data on *An. gambiae* mosquitoes from the Ag1000G project [32]. The Ag1000G dataset includes sequences from 16 mainland African populations and from populations on Mayotte and Bioko, two islands 500km off the east and 30km off the west coast of Africa, respectively. Note that in presenting this analysis we are not advocating the use of double drives on these islands, and merely wish to investigate whether the requisite differentiation can be found on island populations. For our analysis we focussed on potential PAM sequences (NGG or CCN), on the logic that a construct would be unlikely to mutate to recognise a new PAM, whereas this could occur for a protospacer. The entire dataset includes 57 million polymorphic sites, which we screened for PAM sites present in the island population and at a frequency <10%, <5%, or absent from all other populations. In Mayotte, for PAM sequences that were completely private to the island (i.e., not found in any other population), only 1 of them had no pre-existing resistance (i.e., was found in all 48 sequences from the island), whereas 25 had pre-existing resistance less than 20%, and 353 had pre-existing resistance less than 50%. PAM sequences with small but nonzero frequencies on the mainland were even more abundant (Fig 7). Bioko island is not as differentiated as Mayotte from the mainland populations, and the sample size is smaller (18 sequences), but still there are some potential candidate sites.

**Fig 7.**
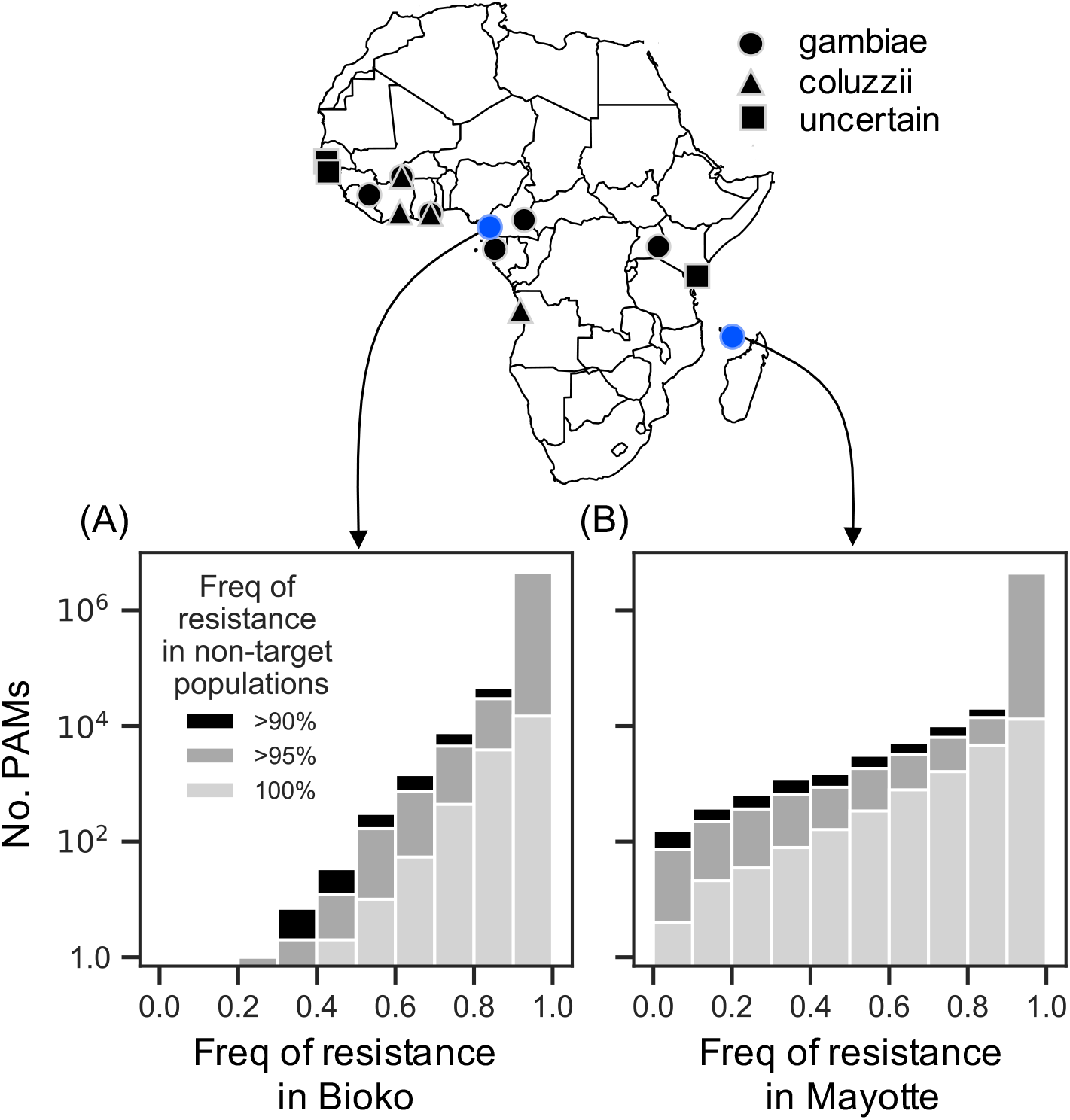
Frequency of PAM sites in island populations of An. gambiae. Numbers of PAM sites (NGG or CCN) with varying frequencies of resistance in samples of *An. gambiae* from two oceanic islands (blue map labels): **(A)** Bioko island (n = 18 sequences, from 9 individuals), and **(B)** Mayotte island (n = 48 sequences, from 24 individuals), where the PAM site frequency in each non-target population (black map labels) is <10%, <5% or 0% (i.e., target site resistance is >90%, >95% or 100%). GG or CC dinucleotides which varied by at least one base were considered to be resistant.

## Discussion

Given that some of the most promising gene drive approaches for population control use (CRISPR-based) sequence-specific nucleases, an obvious way to limit their spread and impact is to exploit sequence differences between target and non-target populations. In this paper we have proposed using a double drive design, here defined as one that uses two constructs, inserted at different locations in the genome, both of which can increase in frequency, and which interact such that the transmission of at least one of them depends on the other. Previously published examples that fit this definition include those for 2-locus under-dominance [14, 15, 33, 34], and Medusa [35], tethered [36], integral [26], and transcomplementing [37] gene drives. As with single-construct gene drives, these various proposed designs differ in purpose (suppression vs. modification), release rate needed to initiate spread (low vs high threshold), and the molecular basis for the superMendelian inheritance (homing, toxin-antidote interactions, or a combination of the two). The requirement that both constructs can increase in frequency over time excludes split drives [25, 38–40] and killer-rescue systems [41, 42], in which only one of the two components increases in frequency. In our proposed designs there is a division of labour between the two constructs, with one responsible for the desired impact (suppression or replacement) and the other for the population restriction, such that together they act as a double drive in the target population and as a split drive in non-target populations. Note that if there are multiple populations of the same species requiring control, each with a different private allele, the same α construct could be used in each case, with only a change in the insertion site of the β construct and the corresponding gRNA. This flexibility may be particularly useful when the α construct requires significant optimisation [26].

We have considered a range of double drive designs of increasing resilience, as judged by their ability to cope with an increasing frequency of pre-existing resistance at the differentiated locus. The simplest designs do not have any component beyond those needed for any CRISPR-based construct, and so should be widely applicable [37]. More powerful constructs can be made by adding an X-shredding sex ratio distorter to the load-inducing construct; these have been most effectively demonstrated in *An. gambiae* mosquitoes [11, 43], but may also work more broadly [44]. In other species there are other ways to distort the sex ratio [45–47], and it would be interesting to model whether these alternatives would be expected to have the same impact as an X-shredder in the context of a double drive. An even simpler way to increase the load would be to add extra gRNAs to the α construct that mutate other female fertility genes [48, 49]. The most powerful design we considered targets a female-specific haplo-insufficient gene, or otherwise causes dominant female sterility or lethality. Such genes are not common, but there are some possible candidates [50–52], and our modelling motivates the search for others. Finally, performance (in terms of being able to cope with ever higher frequencies of pre-existing resistance) could presumably also be improved by using a third construct, to construct a triple drive, though modelling would be required to explore the implications of the many different configurations this extension would allow.

The proposed strategy requires that there be a differentiated locus between target and non-target populations. It need not be an essential gene, and could even be selectively neutral. Our focus has been on using so-called private alleles – sequences that are present (but not necessarily fixed) in the target population, and absent (or of negligible frequency) in non-target populations. Our analysis of PAM sites in *An. gambiae* indicates that appropriately differentiated sites may exist in island populations of this species, though our analysis must be considered preliminary: the dataset does not include mainland sites in closest proximity to the island populations, where differentiation may be lower, and we have not considered potential polymorphism in the protospacer sequence (which, if present, may require the use of multiple gRNAs). We have focussed on nucleotide variation at PAM sites on the assumption that a construct is unlikely to mutate to recognise a new PAM; structural variation in the protospacer region may also be an appropriate basis for geographically restricting double drive spread. We have also not attempted to determine whether the observed differentiation is due solely to mutation and drift, or if selection may be involved as well.

Note that the single drives modelled by Sudweeks et al. [19] require the opposite type of differentiation: sequences that are fixed in the target population, even if not private (i.e., even if found at appreciable frequencies in the non-target population). In this latter scenario the challenge is not so much to have an impact on the target population as to not have an impact on the non-target population. What constitutes “acceptable non-impact” may differ widely from one use case to another and must be assessed on a case-by-case basis: in some circumstances spread of the construct and a transient decline in population size followed by recovery may be acceptable, whereas in others any significant spread of the construct may be unacceptable, regardless of impact on population size. Designs with non-autonomous homing of the β construct (Designs 2, 4, and 5) should be less likely to increase in frequency in the non-target population, and may therefore be preferable. We have focused in this paper on differentiated loci on autosomes, but note that for Design 5 the X-shredder is required for the spread of the α construct and, in principle, one could achieve population-restricted spread if the shredder targeted a population-specific sequence on the X chromosome (rather than inserting it into a population-specific autosomal sequence). In many species the X chromosome shows greater population differentiation than autosomes [53], so this alternative may be useful. Finally, if there are no private alleles in the target population, it may be worthwhile considering a two-step approach of first introducing a private allele into a population and then using that allele to control the population [18]. The ability of double drives to exploit private alleles that are selectively neutral and that have a frequency of only 50% (suppression) or 20% (modification) potentially makes this approach more feasible than would otherwise be the case.

In this paper we have used a simple high-level modelling framework in which the generations are discrete, the population is well mixed, and dynamics are deterministic. This framework is appropriate for strategic models aiming to identify candidate approaches that are worthy of further investigation. For any specific use case the appropriate tactical models would need to be developed that incorporate more biological detail, including spatial and stochastic effects. Such extensions will be particularly important when the goal is to eliminate the target population, which is not possible in our deterministic models. Instead, we have reported the minimum relative population size achieved, which is expected to be related to the size of a population that could be eliminated, but determining the precise connection will require bespoke modelling tailored to a specific situation. Further extensions would be needed to allow for on-going movement between target and non-target populations – if there is on-going immigration into the target population, and this cannot be stopped, then it may not be possible to eliminate the target population with a single release of a double drive. Nevertheless, such a release may be sufficient to suppress the population to such an extent that it can be controlled by other means, including recurrent releases of the same constructs. If one is able to achieve an initial release rate of 1% into a target population, and that suppresses the population by a factor of 1000, then the same releases going forwards will constitute a 10-fold inundation, and self-limiting genetic approaches may be sufficient.

## Methods

The basic modelling structure follows that of Burt & Deredec [10]. In brief, populations have discrete generations, mating is random, there are two life stages (juveniles and adults), and juvenile survival is density dependent according to the Beverton-Holt model, which has two parameters, but since we report results in terms of relative population sizes, only one matters, the intrinsic rate of increase (R_m_). We assume this is equal to 6 [49]. Genetic parameter values (rates of DNA cleavage, rates of alternative repair pathways, and the sex ratio produced by X-shredding) are as estimated from *An. gambiae* [11, 21, 54]. Constructs may be inserted into a haplo-sufficient or haplo-insufficient female-essential gene (in which case gene function is disrupted), a selectively neutral sequence (in which case the insertion is also selectively neutral), or a haplo-sufficient or haplo-insufficient gene required for male and female viability (in which case the insertion is again selectively neutral, because it contains a re-coded version of the target gene [23–25], or is inserted in an artificial intron [26]). For constructs inserted into an essential gene we assume end-joining repair produces non-functional cleavage-resistant alleles [21, 55], while for constructs inserted into selectively neutral sites the products of end-joining repair are also neutral. In all models we assume individuals with an intact CRISPR system suffer a 1% fitness cost for every different gRNA they carry as a cost of off-target cleavage, and for population replacement we assume the cargo gene imposes a 20% fitness cost on females. Both these costs are assumed to be dominant. For simplicity, we assume all fitness costs affect survival after density dependent juvenile mortality and before censusing (e.g., as if pupae die). All results are for populations censused at the adult stage. Releases are of heterozygous adult males at 0.1% of the pre-release number of males, and if the two constructs are linked then they are assumed to be in cis; for constructs released in separate males we assume release rates of 0.05% of each. A list of parameters and their baseline values is given in S1 Table and S2 Table. For the PAM site analysis we screened the Ag1000G phase II SNP data for PAM sites (GG or CC dinucleotides) showing variation between samples at one or both nucleotides. PAM site frequencies were calculated per sampling location and filtered for those present in the island population and at <10%, 5%, or absent from all other populations, excluding those containing >5% missing data in at least one sampled population. Further details are given in the S1 Appendix.

## Acknowledgments

We thank Alistair Miles, Nick Harding and Christopher Clarkson for advice on analysing the Ag1000G data and John Connolly, Silke Fuchs and John Mumford for useful comments on a previous draft.

## Supporting Information

### S1 Text. Supplemental methods

#### Genetics and fitness effects of disrupting host genes

Our baseline model has 2 autosomal loci each with 3 alleles. At the first locus A is the wildtype allele; α is the transgenic construct inserted into and disrupting the wild type allele; and a is a cleavage-resistant allele produced by end-joining repair. For population suppression we assume that the A locus is needed for female survival past the pupal stage, that the resistant allele is non-functional, and that functional resistance is not possible [1]. When A is a haplo-sufficient gene, we assume no reduction in survival probability when carrying only one functional copy of the gene. At the second locus there are initially 2 segregating alleles of equal fitness, B and b, with B being more common in the target population than in the non-target population (and vice versa for b). The third allele, β, is the transgenic construct which can home into B but not into b. Where the differentiated B locus is neutral, we assume all end-joining repair events produce cleavage-resistant alleles (b). In the case where the differentiated B locus is an essential gene, we extend the model to have a 4th allele at this locus, bj, which is a non-functional, cleavage resistant, version of B that can be formed by end-joining repair and assume all end-joining leads to non-functional alleles. We also assume that the β construct has been engineered to have minimal effects on fitness (e.g., by having a recoded version of the B allele, or being inserted into an artificial intron). For comparison we also model a single drive consisting of a single construct homing into a female viability gene, and allow for both functional resistance genes that may pre-exist in the population and non-functional resistance alleles created by end-joining (4 allele model; Fig 3 in main text).

For population replacement where α carries a desirable cargo gene and β is in the differentiated locus, we consider three cases, where (i) both A and B loci are selectively neutral; (ii) the A locus is selectively neutral, and B is a haplo-insufficient essential gene; or (iii) the A locus is a haplo-insufficient essential gene, and the B locus is selectively neutral. In those cases where the A or B locus is an essential gene, the α or β construct is assumed to have been designed to have minimal effects on fitness (as above), end-joining produces non-functional alleles which are dominant lethal, and any pre-existing resistant allele (at B) is selectively neutral. In each case there are 3 alleles at each locus, except when B is an essential gene, when there are 4. For comparison we also consider a single drive consisting of a construct that homes into a neutral site (3 allele model) or into a haplo-insufficient essential gene (4 allele model; Fig 6 in main text).

Where the fitness of either A/A or B/B homozygotes is standardised to 1, the fitness of individuals homozygous for the construct α/α or β/β is 1 – *s*_*I*_; homozygous for the cleavage resistant allele a/a or b/b is 1 – *s*_*R*_; heterozygous for the construct A/α or B/β is 1 − *h*_*I*_*s*_*I*_; heterozygous for the cleavage resistant allele A/a or B/b is 1 − *h*_*R*_*s*_*R*_ or heterozygous carrying both the construct and cleavage-resistant allele α/a or β/b is 1 – *s*_*IR*_, where *s*_*I*_, *s*_*R*_ and *s*_*IR*_ are selection coefficients and *h*_*I*_ and *h*_*R*_ are dominance coefficients, each of which differ depending on the locus (A and B) and between sexes (see S2 Table for more details of locus-specific fitness costs for each scenario modelled).

#### Construct activity and fitness effects

Each construct consists of one or more transcription units (Cas9, gRNA, X-shredder, Cargo), and the activity associated with each is assumed to be dominant. For individuals carrying at least one Cas9, one gRNA and the site targeted by the gRNA (A or B), cleavage of the target site occurs with probability *c*, after which end-joining repair occurs with probability *j* converting the target site to a resistant allele (a or b), and homing occurs with probability 1 − *j* converting the target site to the construct (α or β). For males carrying at least one X-shredder, sperm is produced carrying Y or X chromosomes at a ratio of *m*: 1 − *m*. In all models, we assume a fitness cost due to off-target cleavage (*sH*) of 1%, where fitness is reduced by a factor 1 − *sH* if there is at least one Cas9 and one gRNA present, or by a factor (1 − *sH*)^2^ if there is at least one Cas9 and two different gRNAs present. To model costs due to somatic expression of the nuclease, individuals carrying at least one Cas9, one gRNA and the site targeted by the gRNA, have fitness reduced by a factor 1 − *sS*. Similarly, to model costs associated with X-shredding, males carrying at least one X-shredder have fitness reduced by a factor 1 − *sX*. With the exception of Fig 5B and C in the main text, we assume both *sS* and *sX* to be zero. For population replacement we assume the cargo has a fitness cost (*sC*) of 20% in females, with no effect on males. In all cases the fitness costs associated with each unit’s activity are assumed to be dominant, and for simplicity, are assumed to affect survival of individuals after density dependence and before censusing (e.g., at the pupal stage).

The overall fitness of an individual is therefore the product of the relative fitness due to host gene disruption at locus A, host gene disruption at locus B and construct activity.

#### Analyses of evolutionary stability

We extend both the 3-allele and 4-allele models to allow for loss-of-function mutations of each transcription unit (Cas9, gRNA, X-shredder, cargo). After release of fully functional constructs, we assume that each unit mutates with probability *μ*; mutation can only occur during homing; it is possible for more than one unit to mutate during a single homing event; and mutation back to a functional unit does not occur. Constructs carrying non-functional units are expected to carry the fitness costs associated with disrupting the function of the host gene, however, no longer carry costs due to the transcription unit activity (e.g., off-target cleavage). Although at the sequence level constructs carrying only non-functional units are expected to differ from cleavage-resistant alleles produced through end-joining repair, both are assumed to be resistant to cleavage and carry the same fitness costs, therefore we model them as a single allele.

#### Population biology

We model a population with discrete, non-overlapping generations. In each generation, we assume the population mates randomly, that females produce *f* fertilised eggs and that males are not limiting in their production. Juvenile survival is density-dependent, such that the probability of surviving is 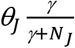, where *θ*_*J*_ juvenile is the density-independent probability the survives to adulthood, *γ* determines the strength of density dependent mortality and *N*_*J*_ is the total number of juveniles in the population during the generation. The intrinsic rate of increase (R_m_) of wild-type population is 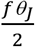. After density-dependent juvenile mortality, additional genotype-dependent mortality occurs before juveniles mature into adults, taking into account fitness costs associated with the constructs and cleavage-resistant alleles. Adult males and females then produce gametes, during which recombination occurs between the A and B loci with probability *r*, and, depending on the genotype, cleavage followed by homing or mutation may occur in males or females, and shredding of the X-chromosome may occur in males.

#### Population censusing

Relative population densities and allele frequencies are of populations censused at the adult stage and the correlation between the constructs α and β is:

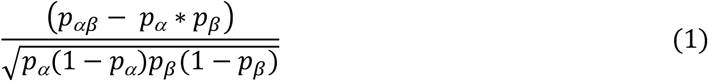

where, *p*_*αβ*_ is the frequency of αβ chromosomes in the population and *p*_*α*_ and *p*_*β*_ are the frequency of α and β alleles in the population respectively, each calculated from populations censused at the adult stage.

#### PAM site analysis in An. gambiae

To identify PAM sites (NGG or CCN) in *An. gambiae* with the type of differentiion required for our approach, SNP variants from the Phase II Ag1000G dataset were first screened for the presence of at least one G or C allele. Positions upstream and downstream of the variant were then screened to calculate frequencies of GG and CC dinucleotides, using the AgamP4 reference genome for invariant neighbouring sites, or the Ag1000G SNP data for neighbouring variants. Since phased data was only available for biallelic variants, dinucleotides containing two variants, of which at least one was multiallelic, were not screened (24% of G or C variants). For each differentiated PAM site, frequencies were calculated for the 16 sampled populations (Cameroon *An. gambiae* n=594, Uganda *An. gambiae* n=224, Burkina Faso *An. gambiae* n=184, Guinea-Bissau uncertain sp. n=182, Angola *An. coluzzii* n=156, Burkina Faso *An. coluzzii* n=150, Cote d’Ivoire *An. coluzzii* n=142, Gabon *An. gambiae* n=138, The Gambia uncertain sp. n=130, Ghana *An. coluzzii* n=110, Kenya uncertain sp. n=96, Guinea *An. gambiae* n=80, Mayotte *An. gambiae* n=48, Ghana *An. gambiae* n=24, Bioko *An. gambiae* n=18 and Guinea *An colluzzi* n=8, where n is the number of sequences and n/2 is the number of individuals sampled). PAM sites containing >5% missing data at any sampling location were removed (0.5% PAMs), leaving a total of 13,462,450 polymorphic PAM sites. The per-site PAM frequencies were used to identify PAM sites present in the island population of interest, and at frequencies <10%, <5% or absent in all other populations, excluding the *An. coluzzii* population from Guinea due to low sample size (n=8, from 4 individuals). Analysis of the Ag1000G data made heavy use of the python package scikit-allel (https://doi.org/10.5281/zenodo.3238280).

---

**S1 Table.** Model parameters and baseline values.

| Parameter | Description | Baseline value |
| --- | --- | --- |
| $R_m$ | Intrinsic rate of population increase | 6 |
| $r$ | Recombination between loci | 0.5 |
| $d$ | Proportion of offspring carrying transgene | 0.976 [1] |
| $u$ | Proportion of non-homed chromosomes which are mutant | 0.392 [2] |
| $e$ | Probability of homing | $2d - 1 = 0.953$ |
| $c$ | Probability of cleavage | $e + (1 - e)u = 0.971$ |
| $j$ | Probability of end joining given cleavage | $\frac{(1 - e)u}{e + (1 - e)u} = 0.019$ |
| $m$ | Proportion of Y-bearing sperm produced by X-shredder males | 0.95 [3] |
| $s_H$ | Cost of off-target cleavage in individuals expressing Cas9 and gRNA (targeting either allele A or B) (dominant) | 0.01 |
| $s_S$ | Female cost of somatic cleavage of wildtype allele by Cas9 and gRNA (targeting either allele A or B) (dominant) | 0 |
| $s_X$ | Male cost of X-shredding (dominant) | 0 |
| $s_C$ | Female cost of cargo expression | 0.2 |
| $p$ | Probability of a functional mutant being produced through end joining | 0 |
| $\mu$ | Probability of each component becoming non-functional during homing | 0 |

**S2 Table.**
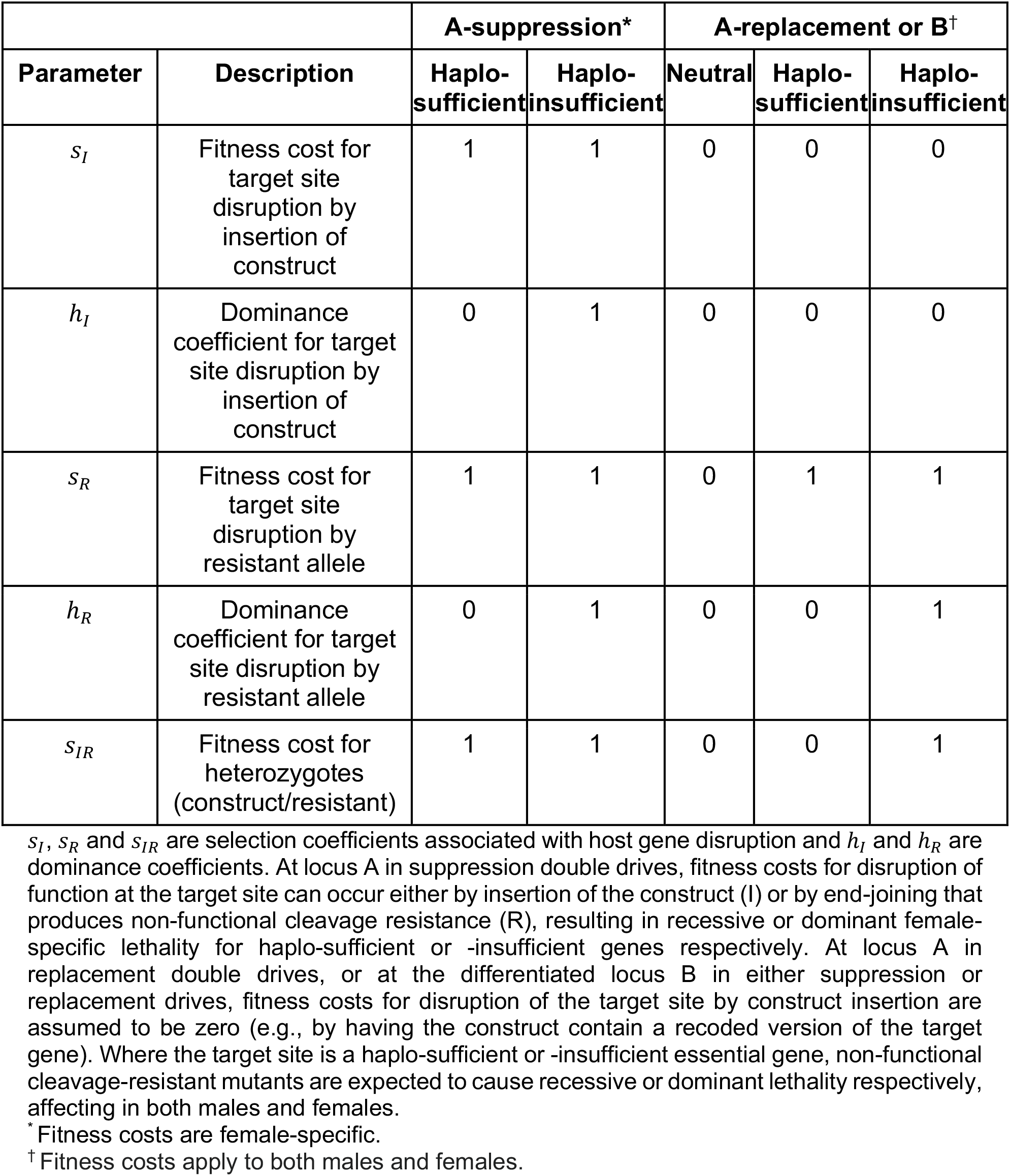
Host gene disruption fitness costs.

**S1 Fig.**
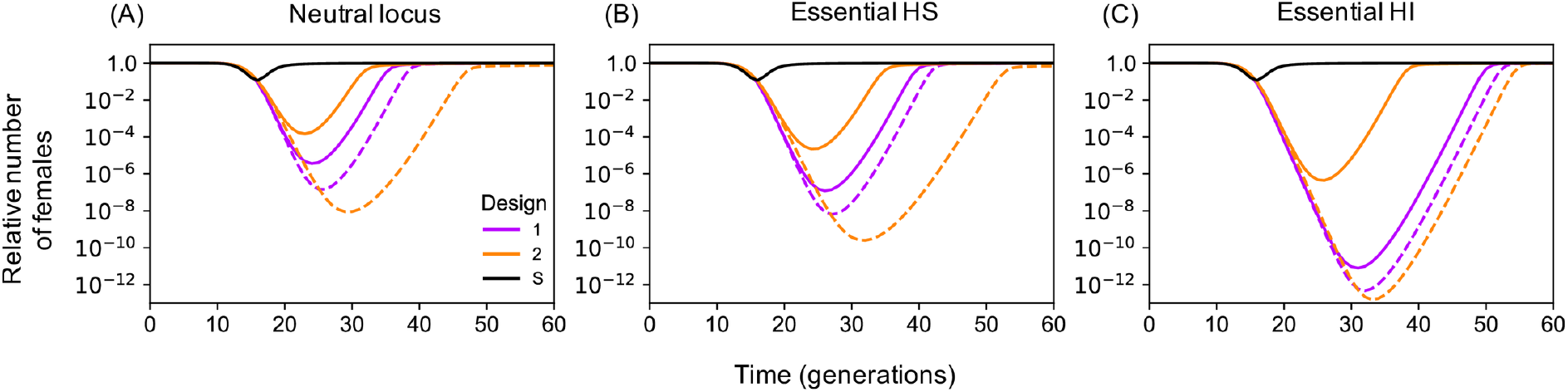
Timecourse for the relative number of females over time for Designs 1 and 2. Solid lines are for where α and β are unlinked and dashed lines for where they are linked (r=0.01). Shown are the cases where β is inserted as a neutral insertion into **(A)** a neutral locus, **(B)** an essential haplo-sufficient gene or **(C)** an essential haplo-insufficient gene. Shown for comparison is a time course for a single drive targeting a haplo-sufficient female-specific viability gene (S).

**S2 Fig.**
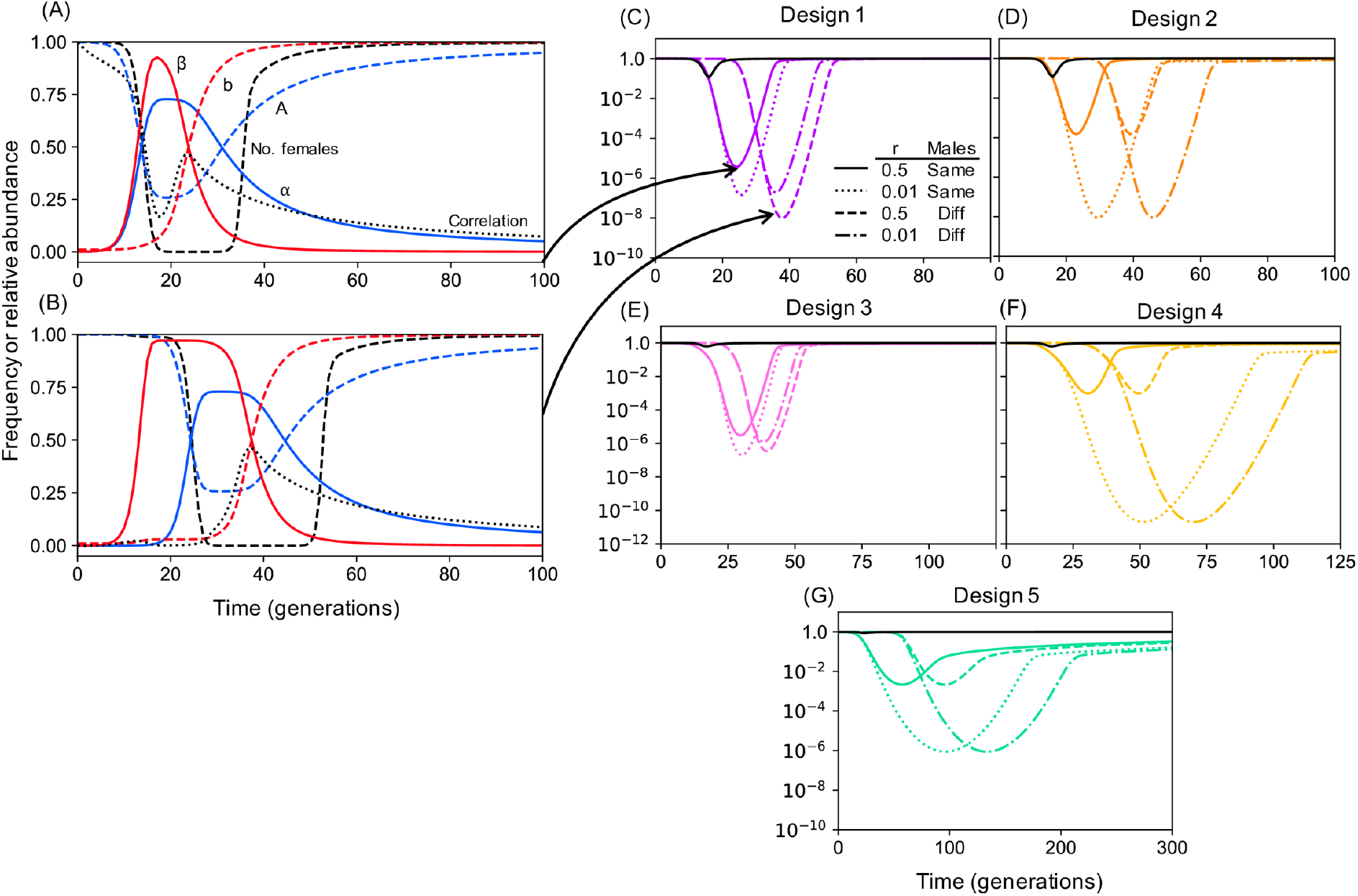
Effect of releasing constructs in the same or different males. Comparison of dynamics for Design 1 when the constructs are unlinked and are released in the same **(A)** or in different **(B)** males. If the constructs are released in separate males the initial correlation between α and β (black dotted line) is negative, allowing β (solid red line) to increase to a higher frequency than if released in the same males as α where it experiences higher fitness costs. Consequently α (solid blue line) is retained at high frequency in the population for longer resulting in a greater reduction in relative number of females, though there is a longer delay between release and impact. **(C-G)** Timecourse for the relative number of females over time for Designs 1-5 where constructs are unlinked and released in the same males (solid lines), linked and released in the same males (dotted lined), unlinked and released in different males (dashed lines) or linked and released in different males (dot-dashed). Pre-existing resistance is assumed to be 1% **(C, D)**, 20% **(E, F)** and 50% **(G)**. For Designs 2, 4 and 5 **(D, F, G)** separate releases only delay the impact because β cannot increase in frequency autonomously, whereas for Designs 1 and 3 separate releases can give a larger (though still delayed) impact when constructs are unlinked, but not when they are closely linked. Shown for comparison is a time course for a single drive targeting a female-specific viability gene with the same level of pre-existing resistance (1%, 20%, or 50%; black solid lines).

**S3 Fig.**
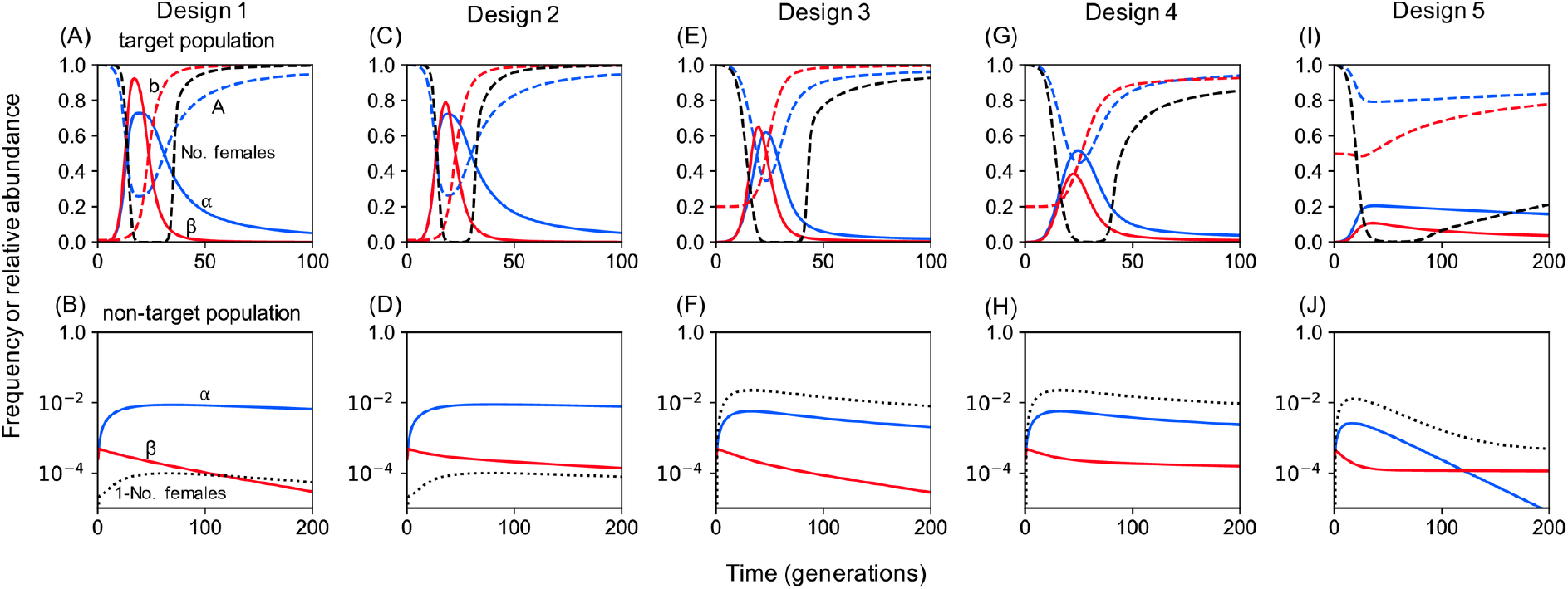
Example time courses for double drives for population suppression. Design 1 **(A, B)** and 2 **(B, C)** assuming 1% and 100% pre-existing resistance in target and non-target populations. Design 3 **(E, F)** and 4 **(G, H)** assuming 20% and 100% pre-existing resistance in target and non-target populations. Design 5 **(I, J)** assuming 50% and 100% pre-existing resistance in target and non-target populations respectively. Plots for Designs 1, 3, and 5 are the same as in the main text, and presented here to facilitate comparisons.

**S4 Fig.**
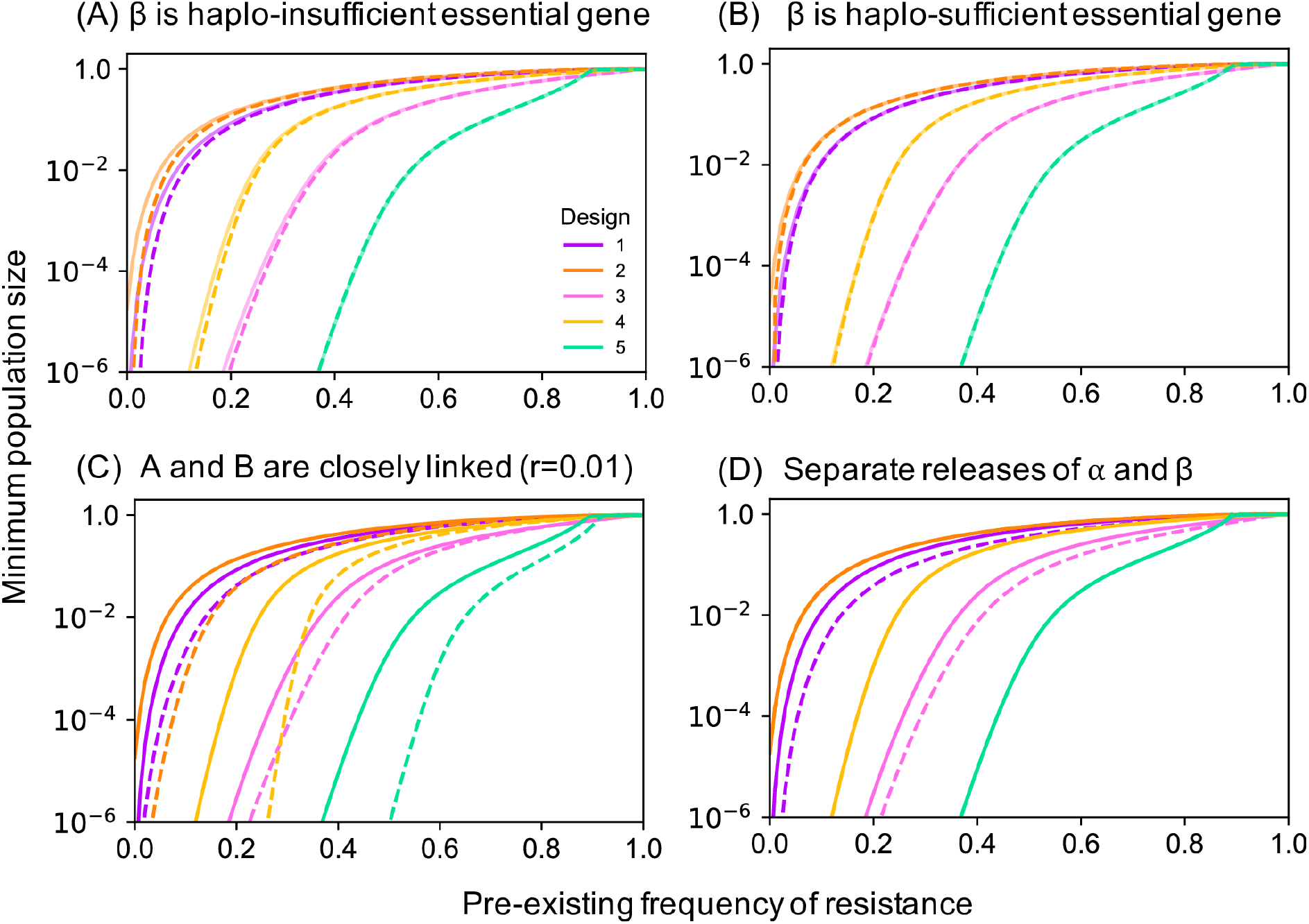
The impact of alternative designs and their variants on population suppression as a function of the pre-existing frequency of resistance. Solid lines are for baseline conditions (α and β are released in the same males, β is in a neutral locus, and loci are unlinked), and are the same in each panel. Dashed lines are for variants where **(A)** β is inserted as a neutral insertion into an essential haplo-insufficient gene, **(B)** β is inserted as a neutral insertion into an essential haplo-sufficient gene, **(C)** loci are linked (r=0.01), and **(D)** α and β are released in separate males, holding all other properties at baseline.

**S5 Fig.**
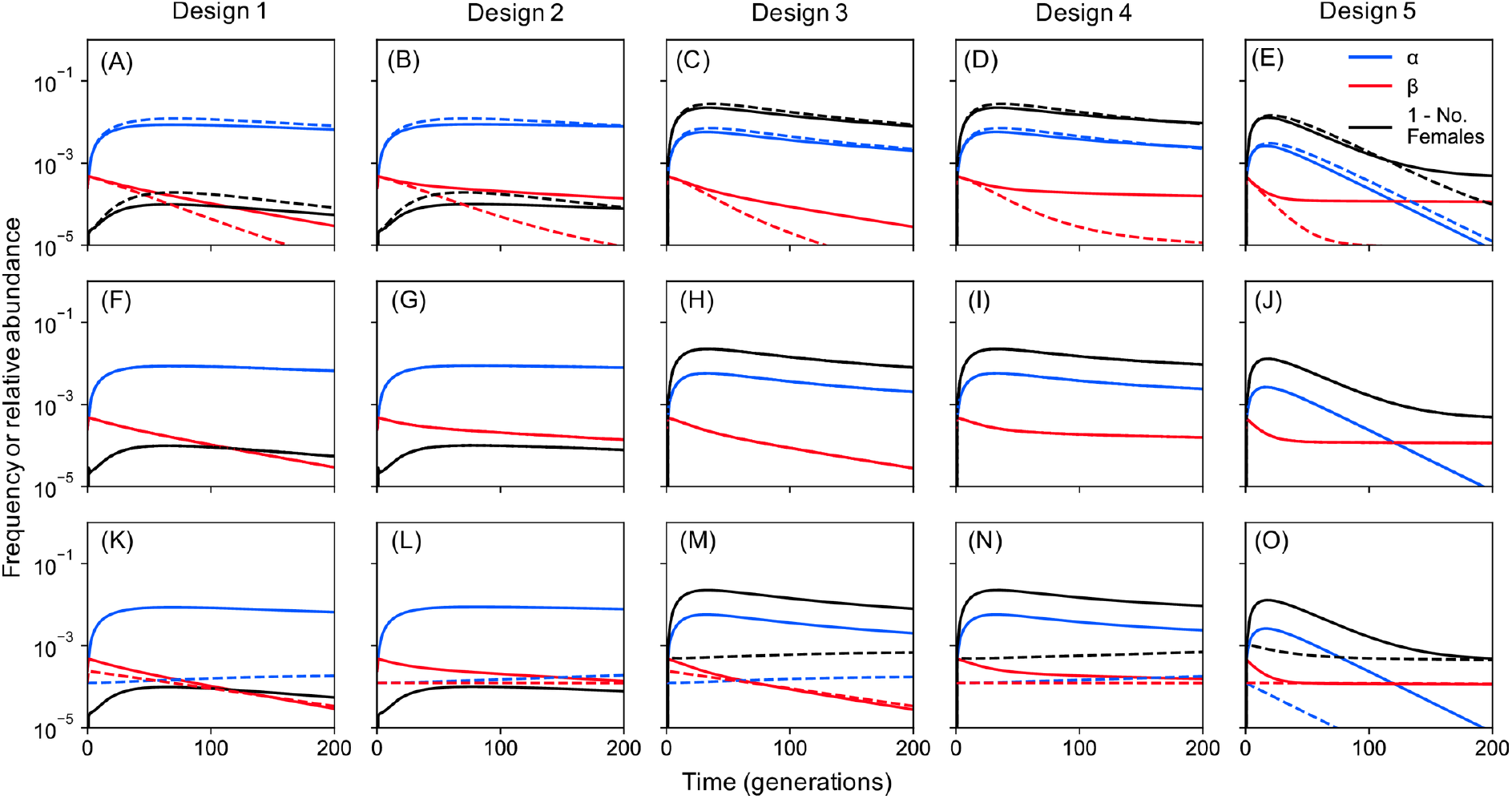
Timecourse of allele frequencies and population suppression (1-relative number of females) for Designs 1-5 in non-target populations where the resistant allele is present at 100%. **(A-E)** α and β are unlinked (solid lines) or linked (dashed lines). **(F-J)** β is inserted as a neutral insertion into a neutral site (solid lines), haplo-sufficient essential gene (dashed lines) or haplo-insufficient essential gene. **(K-O)** α and β are released in the same males (solid lines) or different males (dashed lines).

**S6 Fig.**
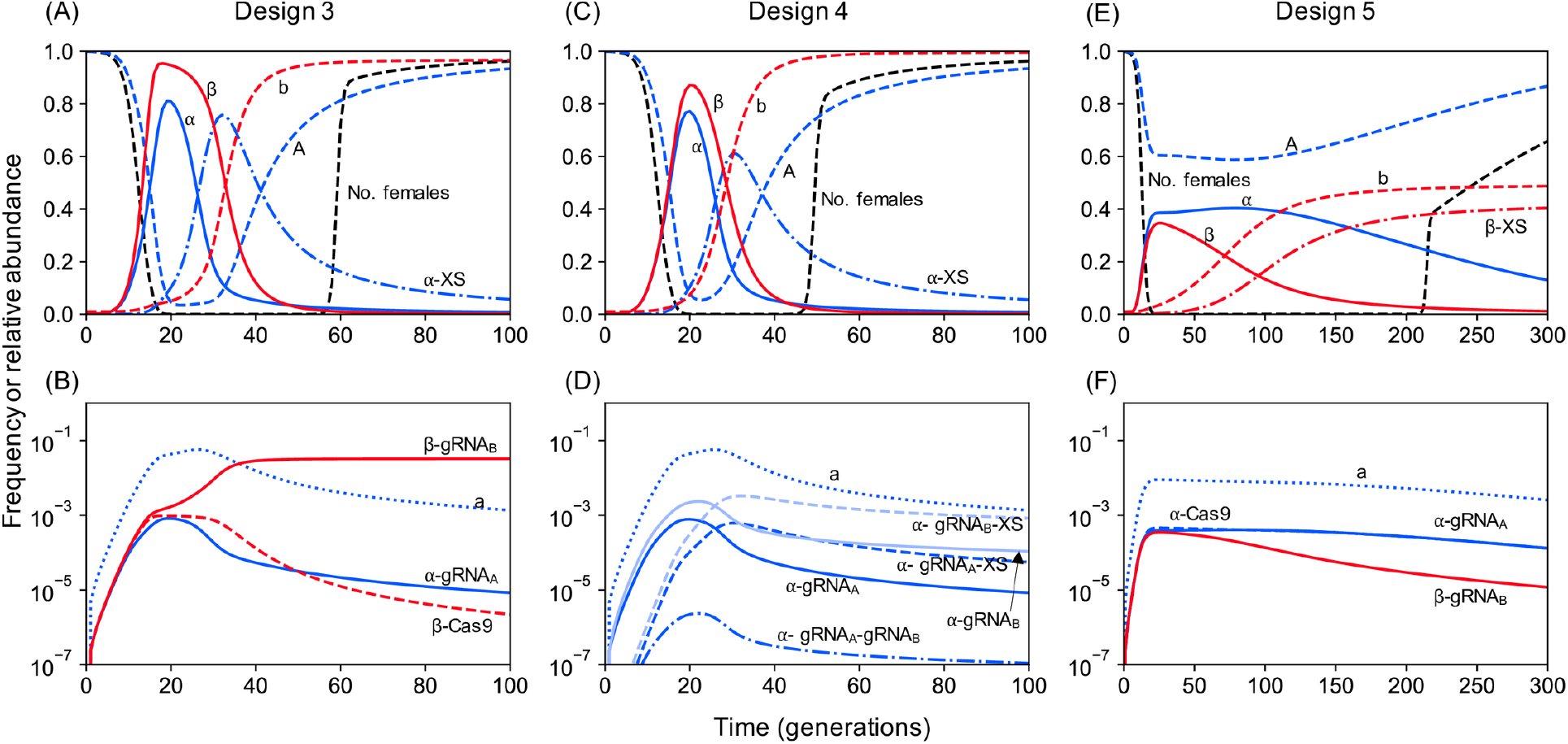
Timecourses for Designs 3, 4 and 5 assuming pre-existing frequency of resistance of 1%, where homing-associated loss-of-function mutations occur for each component of each construct with probability 10e-3. For each design, the intact constructs (α, blue solid lines and β, red solid lines) increase in frequency together, causing the relative number of females (black dashed lines) to decline. For designs 3 and 4, loss-of-function mutations at the X-shredder (α-XS, blue dashed-dotted lines) are selected for, replacing α **(A, C)**. Since α-XS is identical to α in designs 1 and 2, the construct continues to reduce the relative number of females. If the population is not eliminated, β is eventually replaced by the resistant b allele and α-XS is replaced by the wild-type A allele, allowing the population to recover. For design 5, loss-of-function mutations at the X-shredder (β-XS, red dashed-dotted lines) are also selected for, but increases in frequency more slowly than the α-XS allele in Designs 3 and 4, and the intact β construct persists longer. For all designs, loss-of-function mutations at each of the other components (Cas-9, gRNA_A_ and gRNA_B_) remain at low frequency, having negligible impact on the efficacy of the designs **(B, D, F)**.

**S7 Fig.**
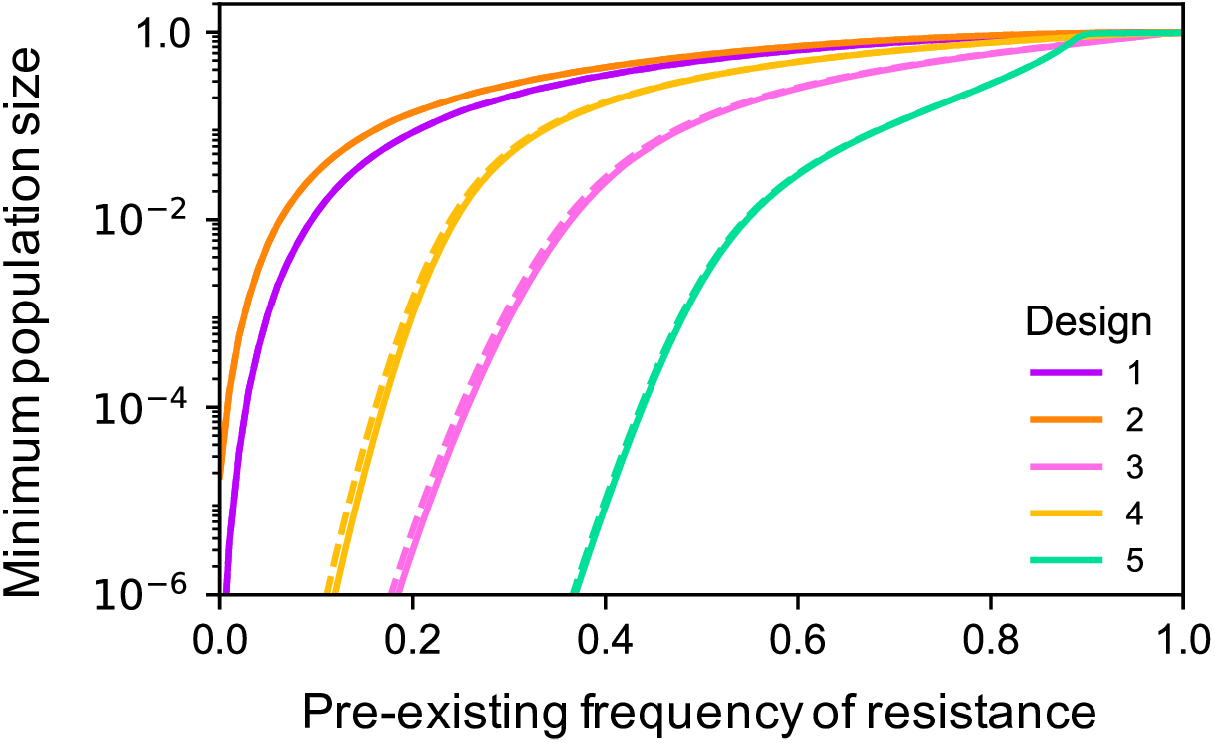
Loss-of-function mutation rates of 10e-4 have minimal impact on the extent of population suppression. Solid lines are for baseline conditions where constructs remain intact after release, while dashed lines are for homing-associated loss-of-function mutations occurring at each component of each construct with probability 10e-4.

